# Large-scale column-free purification of bovine F-ATP synthase

**DOI:** 10.1101/2023.06.01.543336

**Authors:** Jiko Chimari, Yukio Morimoto, Tomitake Tsukihara, Christoph Gerle

## Abstract

Mammalian F-ATP synthase is central to mitochondrial bioenergetics and is present in the inner mitochondrial membrane in a dynamic oligomeric state of higher oligomers, tetramers, dimers and monomers. In vitro investigations of mammalian F-ATP synthase are often limited by the ability to purify the oligomeric forms present in vivo at sufficient quantity, stability and purity. We developed a purification approach for the isolation of bovine F-ATP synthase from heart muscle mitochondria that uses a combination of buffer conditions favouring IF1 binding and sucrose density gradient ultracentrifugation to yield stable complexes at high purity in the milligram range. By tuning the GDN to LMNG ratio in a final gradient, fractions that are either enriched in tetrameric or monomeric F-ATP synthase can be obtained. It is expected that this large-scale column-free purification strategy broadens the spectrum of in vitro investigation on mammalian F-ATP synthase.

## Introduction

Mitochondrial F-ATP synthase is an inner membrane bound rotary nano-machine essential for the interconversion of proton motive force (pmf) and ATP(1, 2). Unlike its bacterial and chloroplast counterparts mitochondrial F-ATP synthase is present in diverse, species dependent oligomeric forms and posesses likewise diverse and a species dependent set of supernumerary subunits that are not directly involved in ATP synthesis(3–12). In mammals at times of a pmf insufficient for ATP synthesis futile ATP hydrolysis is avoided via the binding of the natural inhibitor protein Inhibitor Factor 1 (IF1)(13–16). Binding of IF1 to the hydrophilic F1 domain is thought to stabilize oligomerization(14), a notion that has been greatly strengthened by the structure of the IF1 bound porcine F-ATP synthase tetramer in which two neighboring dimers are connected through two interlinking IF1 dimers(11). A recent study on mammalian F-ATP synthase as a drugable complex for the alleviation of mitochondrial disorder highlighted the usefulness of purified F-ATP synthase for in vitro studies in drug discovery(17). Likewise, the newly discovered role of mammalian F-ATP synthase as the molecular identity of the mitochondrial permeability transition pore (PTP) greatly raises interest in the ability to conduct conclusive in vitro experiments using purified F-ATP synthase complexes in their various oligomeric forms(18–20). The above underlines the need to develop novel protocols that ease the purification and experimental handling of mammalian F-ATP synthase in both monomeric and oligomeric states.

Since its discovery of being the site of ATP synthesis from ADP and Pi more than 50 years ago(21, 22), mammalian F-ATP synthase isolated from heart muscle tissue mitochondria has been the subject of investigation on the structure and function of mitochondrial F-ATP synthase. Concomitantly, various strategies for the isolation of mitochondrial F-ATP synthase from mammalian cells were developed(23–29).

Early studies succeeded to purify its water soluble F1 domain to high purity, eventually allowing the growth of well diffracting 3D crystals for structure determination by X-ray crystallography(30, 31). Isolation of the whole mammalian F-ATP synthase complex, however, proved to be challenging when using conventional detergents, often yielding only complexes of poor stability and homogeneity. Recently, the use of very mild, lipid like detergents has dramatically improved the situation in regard to the stability of mammalian F-ATP synthase during isolation of the complex. Namely, the natural compound digitonin, its synthetic stand-in GDN(32) and the two acyl chain tailed LMNG(33) have become the detergents of choice for isolation of intact F-ATP synthase from mammalian source. All of them greatly ease the instability problem and enabled the successful large-scale purification of F-ATP synthase from heart muscle tissue of large animals or human cell culture for structural and functional studies(9–11, 17, 19, 34–38). However, isolation of highly stable monomeric and especially oligomeric mammalian F-ATP synthase at high yield and purity is still challenging. IF1 bound tetrameric F-ATP synthase is perhaps the most desirable of all oligomeric forms, since it represents a minimal oligomeric structural unit which can be expected to be of high physiological relevance.

In all of the reported large-scale purifications of mitochondrial F-ATP synthase from mammalian cells, chromatography steps such as anion exchange chromatography, size exclusion chromatography or IF1 affinity resins are employed as necessary tools for the removal of contaminating proteins(10, 11, 19, 39–41). In contrast, it has been shown that for large membrane protein complexes from higher organisms isolation procedures that do not involve any chromatography steps can be beneficial for obtaining well behaving protein suitable for structural and functional studies(42–44).

We reasoned that in vitro studies of mammalian F-ATP synthase could greatly benefit, if novel strategies for large-scale column-free purification would be developed. This might enable design of purification strategies tailored to the isolation of specific oligomeric states while avoiding any damage during the purification process. Density gradient ultracentrifugation is a powerful method for column-free, mild isolation of large protein complexes widely used in photosynthetic and ribosomal research fields(43, 45). The size similarity of monomeric, dimeric and tetrameric F-ATP synthase to other complexes of the mitochondrial respiratory chain, however, complicates application of this approach to mammalian F-ATP synthase. Given that higher oligomers of F-ATP synthase are much larger than respiratory supercomplexes, and that IF1 binding can stabilize oligomer formation, we surmised that a combination of IF1 binding and density gradient ultracentrifugation might be a possible avenue for developing novel procedures for large-scale column-free purification.

Here, we describe the successful large-scale column-free purification of oligomeric and monomeric F-ATP synthase from bovine heart mitochondria at high yield and purity by sucrose density gradient ultracentrifugation.

## Experimental Procedures

### Isolation of inner mitochondrial membrane

The procedure for the isolation of mitochondrial inner membrane employed here is basically identical to the one used for 3D crystallization of bovine cytochrome c oxidase as reported previously(28). Slight modifications, however, were made to buffer composition and pH to stabilize IF1 binding. In brief, a single bovine heart was obtained from a local abattoir immediately after animal sacrifice. Fat tissue was removed and 600 g of lean, red meat minced using a commercial meat mincer. The mince is placed into 2.3 liter of ice-cold distilled water supplied with 350 ml of ice-cold phosphate buffer (0.2 M NaPi, pH 7.4), and a spate of PMSF. This mixture is homogenised in a Polytron PT3100D homogenizer at 11,000 rpm for 10 minutes. Larger cell debris is removed by spin-down at 2,800 rpm for 20 minutes at 4 degree Celsius with a large-scale refrigerated centrifuge (Hitachi Himac CR20G) using a R2A rotor. Subsequently the supernatant is carefully separated from the soft pellet by straining through two sheets of gauze. Thereafter, mitochondria are pelleted from the strained supernatant by centrifugation at 8,000 rpm for 25 minutes at 4 degree celsius with a large-scale refrigerated centrifuge (Hitachi Himac CR20G) using a R12A F rotor and the supernatant discarded. The mitochondrial precipitate is suspended in a buffer of 40 mM HEPES-NaOH (pH 7.3), 5 mM MgCl2, 5 mM DTT, 5 mM EGTA, 0.5 mM ADP and homogenized using a loosely fitting Dounce glass homogenizer by ∼12 up-down movements. From the resulting homogenate the inner mitochondrial membrane fraction is pelleted by ultracentrifugation at 100,000 g using a Hitachi Himac CP80WX ultracentrifuge with a P45A T-angle rotor. After discarding the resulting supernatant by aspiration and careful wiping of residual oil from the ultracentrifugal tubes, the total amount of inner mitochondrial membrane is weighted for later adjustment of the membrane-to-detergent ratio.

### Solubilization of inner mitochondrial membrane

Pellets of inner mitochondrial membrane are suspended in a buffer of 40 mM HEPES-NaOH (pH 7.3), 5 mM MgCl2, 5 mM DTT, 5 mM EGTA, 0.5 mM ADP at a volume to weight ratio of one liter to 560 g. The suspension is homogenized by approximately 12 up-down movements in a tightly fitting glass Dounce homogenizer to obtain inside-out vesicles. Note that the resulting pH of the homogenate will be slightly acidic. Solubilization of the inside-out membrane fraction is performed under constant magnetic stirrer mixing at ice-cold temperature. First, sodium deoxycholate is added to a final concentration of 0.73% (w/v) from a 11% stock solution, second, decyl-maltoside (DM) is added to a final concentration of 0.4% (w/v) from a 20% stock solution, after which solid KCl is added to a final concentration of 72 g/l. Finally, when the KCl salt grains are completely dissolved, glyco-diosgenin (GDN) is added to a final concentration of 0.1% (w/v). At this point the solution should have a pH of around 6.0. To remove insolubilized membranes the solution is ultracentrifuged for 40 minutes at 176,000 g using a Hitachi P45A T-angle rotor. The resulting supernatant is strained through four layers of gauze and kept for equilibrium ultracentrifugation.

### Sucrose gradient equilibrium ultracentrifugation

A two step sucrose gradient is prepared in large fixed angle rotor ultracentrifugal tubes using 16 ml of 2.3 M sucrose and 24 ml 1.6 M sucrose solubilized in a buffer of 40 mM HEPES-NaOH (pH 7.3), 5 mM MgCl2, 5 mM DTT, 5 mM EGTA, 0.5 mM ADP, 100 mM KCl, 0.1 % DM (w/v), 0.02% GDN and 0.02% LMNG.

The solubilized inside-out vesicle solution is gently layered onto the two step sucrose gradient and ultracentrifuged to equilibrium at 176,000 g for 42 hours at 4 degree Celsius using a Hitachi P45A T-angle rotor. The resulting gradient is collected in 2 ml fractions from the bottom of each ultracentrifuge tube using a peristaltic pump and fractions are examined for enrichment of F-ATP synthase with low ATP hydrolysis activity employing a combination of SDS-PAGE, CN-PAGE and the Pullman ATP hydrolysis activity assay. Fractions containing a high concentration of F-ATP synthase, but exhibiting relatively low ATP hydrolysis activity are pooled and diluted in a buffer of 40 mM HEPES-NaOH (pH 7.3), 5 mM MgCl2, 5 mM DTT, 5 mM EGTA, 0.5 mM ADP, 100 mM KCl and 0.02% GDN to a final sucrose concentration that is slightly below 0.6 M sucrose as monitored using a PAL-1 pocket refractometer (Atago).

### Sucrose step gradient ultracentrifugation

A step gradient of sucrose solution is layered into large ultracentrifugal tubes from bottom-to-top using 10 ml of 1.6 M, 10 ml of 1.0 and 10 ml of 0.6 M sucrose solution with all solution prepared using a buffer of 40 mM HEPES-NaOH (pH 7.3), 5 mM MgCl2, 5 mM DTT, 5 mM EGTA, 0.5 mM ADP, 80 mM KCl and 0.02% GDN. The pooled and diluted F-ATP synthase fractions are gently layered onto the step gradient and ultracentrifuged at 176,000 g for 20 hours at 4 degree celsius using a Hitachi P45A T-angle rotor. The resulting gradient is collected in 2 ml fractions from the bottom of each ultracentrifuge tube using a peristaltic pump and fractions are examined for enrichment of F-ATP synthase by SDS-PAGE and CN-PAGE. Fractions enriched in F-ATP synthase are pooled and diluted to 19% sucrose as monitored using a PAL-1 pocket refractometer (Atago) in a buffer of 40 mM HEPES-NaOH (pH 7.3), 5 mM MgCl2, 5 mM DTT, 5 mM EGTA, 0.5 mM ADP, 100 mM KCl and 0.02% GDN.

### Continuous gradient ultracentrifugation for tetramer enrichment

Using a Gradient Master 108, Biocomp a continuous 40-20% sucrose gradient is prepared in swing-out rotor tubes (Open-Top Thinwall Ultra-Clear Tube, 25 x 89mm) with a buffer composition of 40 mM HEPES-NaOH (pH 7.3), 5 mM MgCl2, 5 mM DTT, 5 mM EGTA, 0.5 mM ADP, 100 mM KCl and 0.02% GDN. The diluted F-ATP synthase fractions are gently layered onto the gradient and ultracentrifuged at 113,000 g for 20 hours using a swing rotor P28S (Hitachi). The resulting gradient is collected in 2 ml fractions from the bottom of each ultracentrifuge tube using a peristaltic pump and fractions are examined for enrichment of tetrameric F-ATP synthase by SDS-PAGE, CN-PAGE and negative stain electron microscopy.

### Continuous gradient ultracentrifugation for monomer enrichment

Using a Gradient Master 108, Biocomp a continuous 40-20% sucrose gradient is prepared in swing-out rotor tubes (Open-Top Thinwall Ultra-Clear Tube, 25 x 89mm) with a buffer composition of 40 mM HEPES-NaOH (pH 7.3), 5 mM MgCl2, 5 mM DTT, 5 mM EGTA, 0.5 mM ADP, 100 mM KCl and 0.02% LMNG. For de-oligomerization the pooled and diluted F-ATP synthase fractions are supplemented with LMNG to a final concentration of 0.5% (w/v) and incubated for 2 hours. Subsequently, the solution is gently layered onto the gradient and ultracentrifuged at 113,000 g for 20 hours using a swing rotor P28S (Hitachi). The resulting gradient is collected in 2 ml fractions from the bottom of each ultracentrifuge tube using a peristaltic pump and fractions are examined for enrichment of monomeric F-ATP synthase by SDS-PAGE, CN-PAGE and negative stain electron microscopy.

### Concentration and sucrose removal by PEG precipitation

Pooled fraction enriched in tetrameric or monomeric F-ATP synthase are supplemented dropwise with PEG 20,000 to a final concentration of 8% and mixed by inversion until turning cloudy. Subsequently the mixture was spun down at 15,000 rpm for 20 minutes at four degrees (TOMY M X-307). The supernatant was completely discarded, and dissolved in 30 μl of 40 mM HEPES-NaOH (pH 7.3), 5 mM MgCl2, 5 mM DTT, 5 mM EGTA, 0.5 mM ADP, 100 mM KCl and 0.02% GDN or 0.02% LMNG and mixed gently by finger tapping until completely dissolved.

### Negative stain electron microscopy

An aliquot of 3.0 μL of enriched fraction was diluted x100 and applied to glow-discharged (5 mA, 10 s, Eiko) continuous carbon film coated copper grids (Nisshin EM, Tokyo, Japan). Staining was performed by applying 3.0 μL 2% uranyl acetate solution. After incubation for 30 s, staining solution was blotted using filter paper (Whatman #1; Whatman, Little Chalfont, UK) and dried. Speciment was inspected using a H-7650 HITACHI transmission electron microscope at 80 kV acceleration voltage equipped with a 1× 1K Tietz FastScan-F114 CCD camera.

### Denaturing SDS gel electrophoresis

SDS-PAGE was performed using 10-20% continuous e-PAGEL HR (Atto). For each fraction 5 μl was applied and electrophoresis conducted using a denaturing phoresis buffer (3.0 g Tris, 14.4 g Glycine and 1.0 g SDS in one liter MiliQ water). Protein bands were visualized by SimplyBlue SafeStain (Invitrogen). The EzProtein Ladder (Atto) was employed as molecular weight marker.

### Clear Native gel electrophoresis

Native gel electrophoresis was performed using 3-12% Bis-Tris gels from Invitrogen and native running buffers from SERVA.

### Pullman ATP hydrolysis activity assay

To monitor ATP hydrolysis activity of collected gradient fractions an ATP-regenerating enzyme-coupled assay was used(21). The hydrolysis of ATP by the F-ATP synthase was followed by NADH oxidation at 340 nm at 20 degree Celsius in the absence or presence of oligomycin.

### Determination of protein concentration

Protein concentration was measured using a NanoDrop Lite (Thermo Fisher Scientific) by UV absorbance at 240 nm.

## Results

In this study we established a large-scale column-free purification procedure for bovine heart mitochondrial F-ATP synthase (see Fig.1 for a workflow chart). We demonstrate that this is feasible when taking advantage of the very large size and molecular weight of IF1 stabilized oligomers of heart muscle tissue F-ATP synthase. In a first step of this approach we made use of the well established procedures to obtain large amounts of highly pure mitochondrial inner membranes from bovine heart muscle tissue. This is achieved by differential centrifugation of first mitochondria and then inside-out vesicles of the mitochondrial inner membrane. Experimental conditions for this type of mitochondrial inner membranes isolation had originally been established for the high resolution structure determination of bovine cytochrome c oxidase by X-ray crystallography(46). Mass spectrometry demonstrated mitochondrial inner membranes prepared in this way are only minimally contaminated with the outer mitochondrial membrane or other cellular membranes(47). In order to strengthen the binding of IF1 to mitochondrial F-ATP synthase during isolation of the inner membrane fraction buffer conditions were modified to a lowered pH and the presence of Mg ADP(48). These modifications did not affect the high yield of ∼70 g inner membrane fraction typically obtainable from 600 g of lean heart muscle tissue (Sup. Fig. 1A-C). A mixture of deoxy-cholate and decyl-maltoside had been previously established to be efficient for the solubilization of the highly packed mitochondrial inner membranes(28). However, with the aim of stabilising larger oligomers of IF1 bound F-ATP synthase we supplemented this detergent mixture with the additional mixture of GDN and LMNG (Sup.Fig. 1D, E). Both novel detergents are known for their ability to stabilize fragile membrane complexes during membrane solubilization and purification. In combination with a slightly acidic pH and the presence of Mg ADP for tight IF1 binding a stabilization of higher oligomers of F-ATP synthase was expected. Under these conditions it was possible to subject the solubilized inner membranes to a 42 hours equilibrium sucrose density gradient centrifugation and obtain almost colorless fractions, indicative of the absence of other complexes of the respiratory chain (Fig.2 A; Sup.Fig.1F). When examined by denaturing SDS-PAGE strong bands distinct for the α and β subunits of the F-ATP synthase could be discerned (Fig.2 C). Western blots using an antibody specific for the F-ATP synthase β subunit confirmed the presence of the β subunit in all fractions, indicating the spread of F-ATP synthase over the sucrose gradient (Sup.Fig. 3). The wide spread presence of F-ATP synthase presumeably is the result of higher oligomers of various size. Clear native polyacryl gel electrophoresis (CN-PAGE) had been specifically developed to allow native gel electrophoresis of fragile mammlian F-ATP synthase oligomers(49). Therefore, together with SDS-PAGE we additionally used CN-PAGE to monitor each sucrose density gradient fraction. In the almost colorless fractions bands at high molecular weight typical for oligomeric F-ATP synthase could be observed(19). Importantly lower molecular bands stemming from broken complexes and dissociated F1 domains were not observed (Fig.2B). When we probed the ATP hydrolysis activity of the F-ATP synthase enriched fractions we detected little ATP hydrolysis activity, again indicative of absence of dissociated F1 domains (Sup.Fig.2). The high molecular weight of these ATP hydrolysis ‘silenced’ F-ATP synthase fractions suggest the presence of IF1 stabilized higher oligomers of F-ATP synthase. In contrast, we could observe higher ATP hydrolase activity in the upper colored fractions of the gradient, which is suggestive of the presence of F-ATP synthase of smaller oligomeric state and a smaller proportion of IF1 binding. As a result a single equilibrium sucrose density gradient allows under the experimental conditions used here, to separate large oligomers of IF1 bound F-ATP synthase in fractions that already contain very little other contaminants.

**Figure 1:**
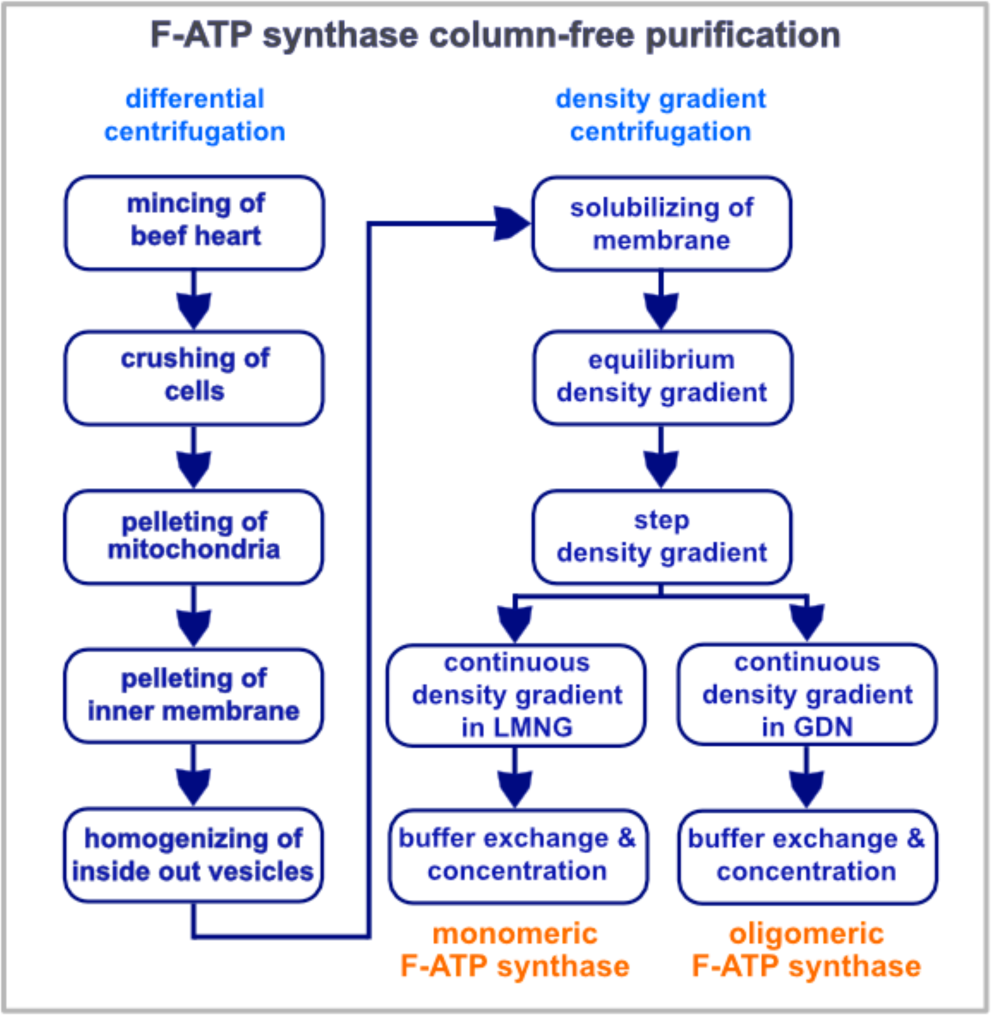
Flow chart of steps in the large-scale column-free purfication of F-ATP synthase from bovine heart muscle tissue mitochondria. Differential centrifugation is employed to obtain large amounts of pure inner mitochondrial membranes. Solubilized membranes are subjected to three sucrose density gradient ultracentrifugation steps which can be tuned via the use of either LMNG or GDN to yield oligomeric or monomeric F-ATP synthase.

**Figure 2:**
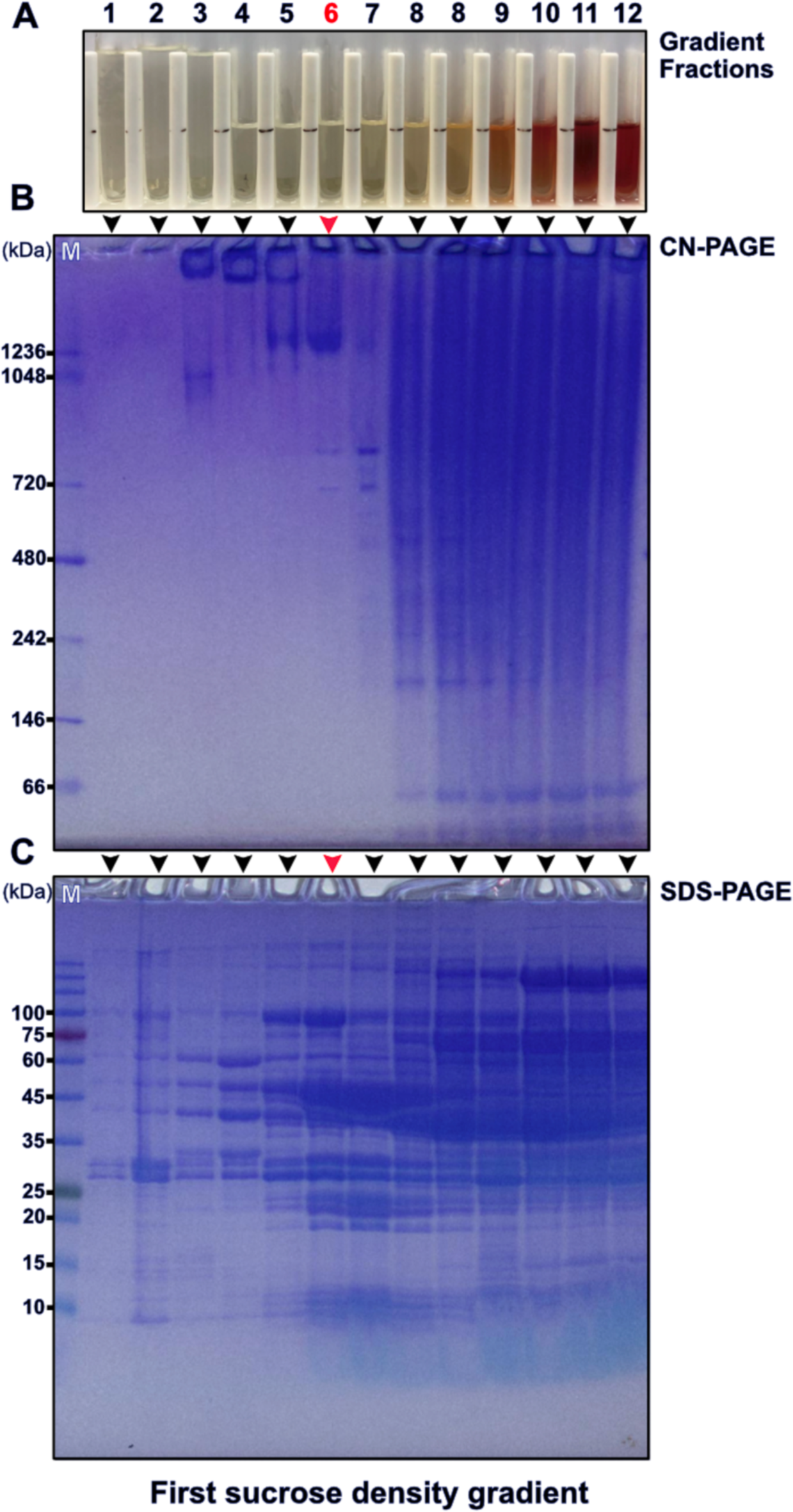
Fractionation of the equilibrium sucrose density gradient. A) Fractions from a gradient (Supplementary Figure 1F, after) as collected from bottom to top. The fraction used for further purification is marked by a 6 in red font. B) Clear native gel electrophoresis of the fractions shown in A) and indicated by arrow heads. The red arrow head indicates the lane corresponding to fraction 6 in A) and M indicating the native molecular weight marker lane. C) Denaturing gel electrophoresis of the fractions as shown in A) and indicated by arrowheads with the red arrowhead marking the fraction used for further purification. A molecular weight marker lane is marked by M.

**Figure 3:**
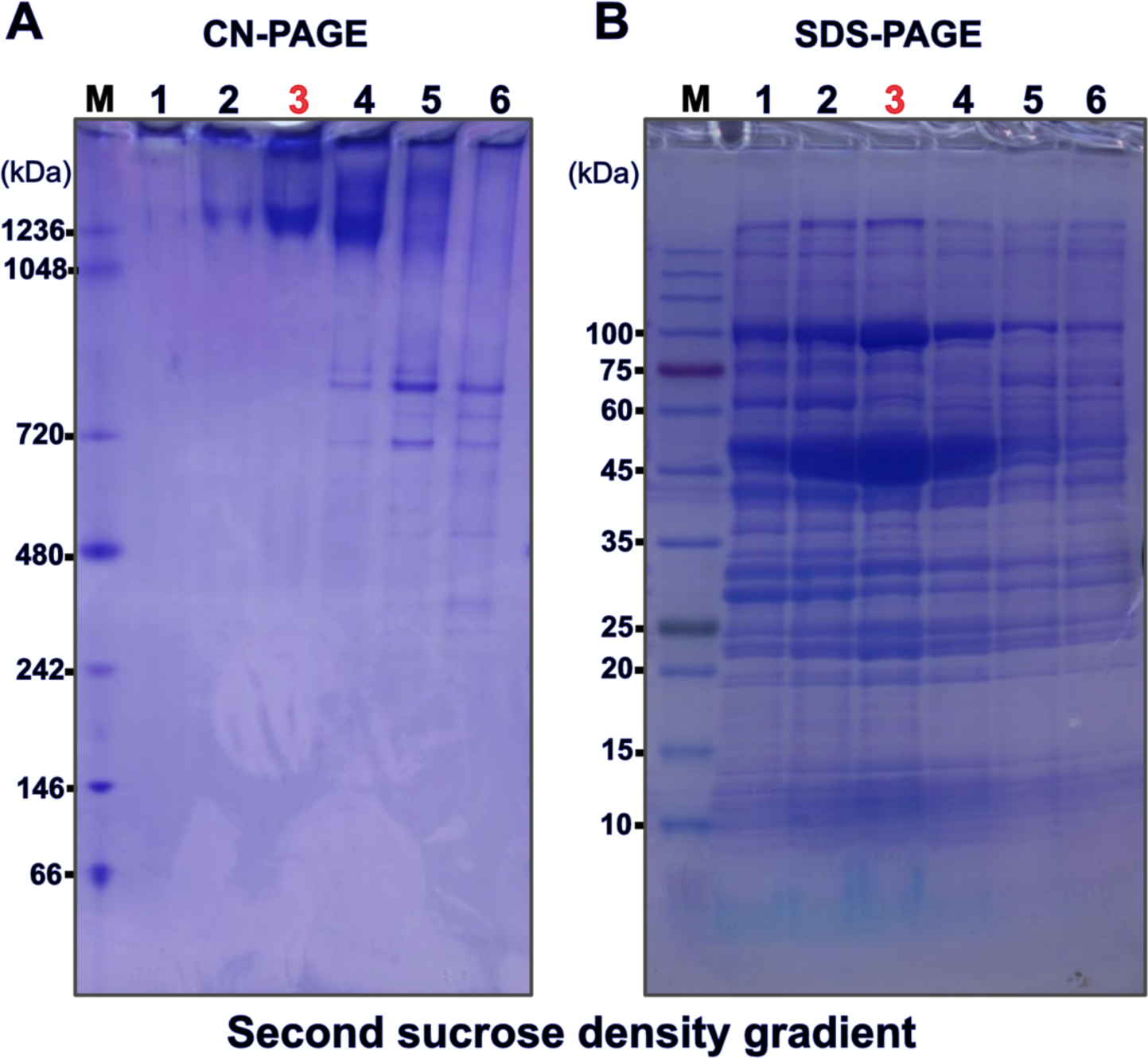
Native and denaturing gel electrophoresis of the second sucrose step density gradient. (Supplemental Fig. 1G, after). A) Fractions from bottom to top as analyzed by clear native gel electrophoresis with the fraction used for further purification indicated by 3 in red font. The molecular weight marker lane on the far left is marked by ‘M’. B) Denaturing gel electrophoresis of the same fractions that were used in A) with a 3 in red font indication the fraction used for further purification. The molecular weight marker lane on the far left is marked by ‘M’.

Next, these colorless, ‘ATP hydrolase activity silent’ F-ATP synthase enriched fractions were pooled and used for a second sucrose step gradient ultracentrifugation for the removal of almost all remaining contaminants. This second sucrose density gradient yielded relatively pure, but oligomeric mixed F-ATP synthase. CN-PAGE of gradient fractions did not show any bands at lower molecular weight indicative of a high stability of the obtained F-ATP synthase fractions also after the second density gradient (Fig.3; Sup.Fig.1G). A typical yield for the indicated target fraction number three (Fig. 3) is about 40 mg of total protein.

A third, continuous sucrose density gradient ultracentrifugation yielded fractions enriched in tetrameric F-ATP synthase, if run in the presence of GDN. Analysis by SDS-PAGE and CN-PAGE of these fractions indicate high purity and stability (Fig.4A,B). As judged by negative stain electron microscopy the proportion of tetrameric F-ATP synthase could be sufficient for structural analysis by single particle cryo-EM (Fig.4C). A typical yield for the ‘tetramer’ fraction is about 10 mg. When supplying LMNG to a final concentration of 0.5% to the F-ATP synthase fractions before running a third, continuous sucrose density gradient in the presence of 0.02% LMNG it was possible to obtain fractions containing almost only monomeric F-ATP synthase. Analysis by SDS-PAGE and CN-PAGE of these fractions indicate high purity and stability (Fig.5A,B). As judged by negative stain electron microscopy the level of purity, stability and monodispersity of the obtained monomeric F-ATP synthase should be sufficient for structural and functional studies (Fig.5C). A typical yield for the ‘monomer’ fraction is about 40 mg. Bovine F-ATP synthase stabilized in LMNG is suitable for rapid formation of proteoliposomes by auto-insertion(38). Therefore we expect the in this way purified monomeric F-ATP synthase could be useful for functional and structural in vitro studies of bovine F-ATP synthase in the context of a proton gradient. Both monomeric and tetrameric F-ATP synthase preparations exhibited long term stability for several days at 4 degree Celsius and if thawed from snap frozen aliquots (Fig.6A,B). Both ‘tetramer’ and ‘monomer’ fractions can be concentrated and buffer exchanged by PEG precipitation for which PEG 20,000 was found to be most suitable among all PEGs tested.

**Figure 4:**
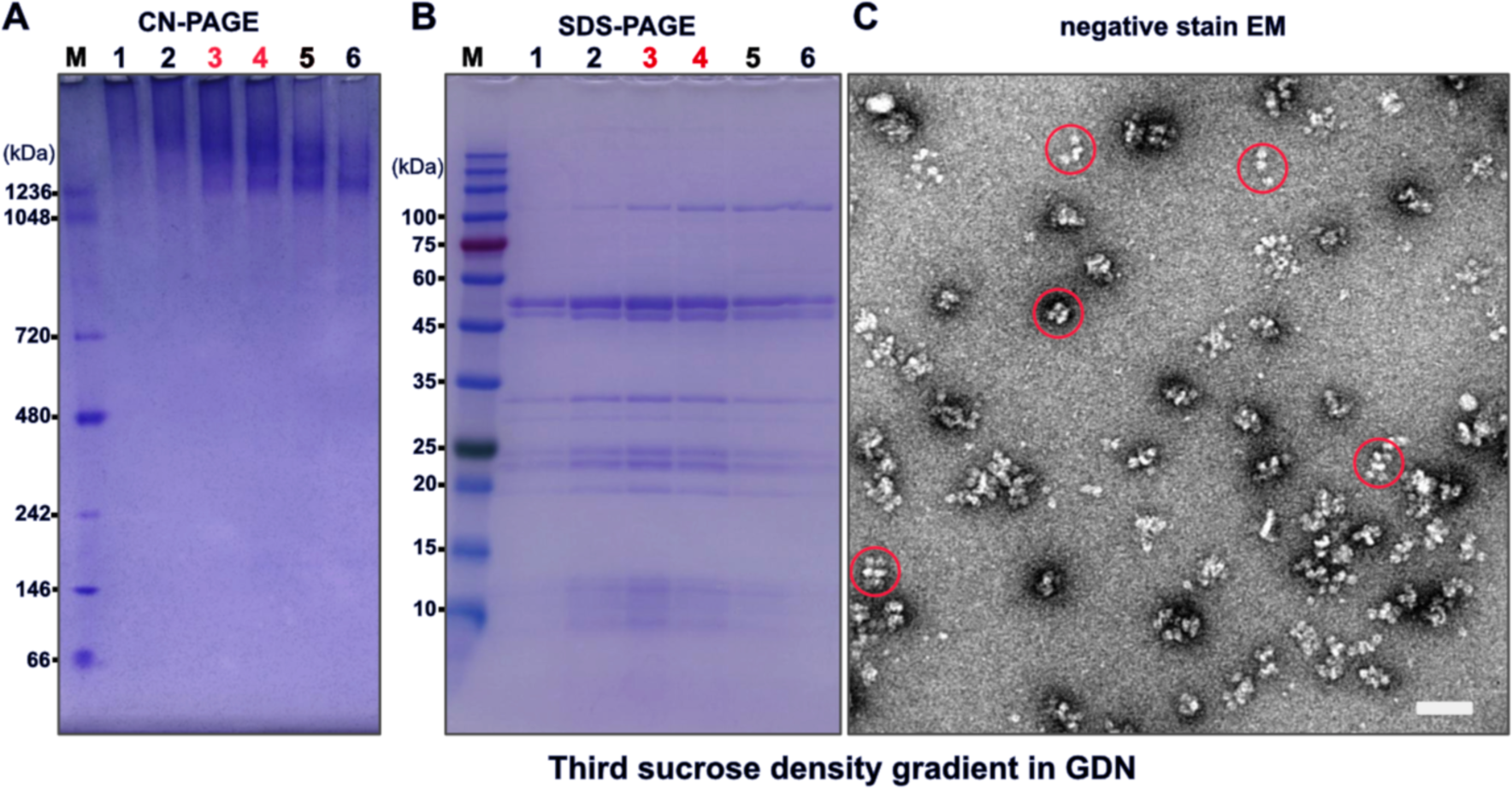
Analysis of the third continuous sucrose GDN density gradient fractions by native gel electrophoresis, denaturing gel electrophoresis and negative stain electron microscopy. A) Clear native gel electrophoresis of fractions collected from bottom to top of the gradient with the fraction enriched in tetrameric F-ATP synthase and used for negative stain electron microscopy marked in red font. The molecular weight marker lane on the far left is marked by ‘M’. B) Denaturing gel electrophoresis of fractions collected from bottom to top of the gradient as analyzed in A). The fraction enriched in tetrameric F-ATP synthase and used for negative stain electron microscopy in marked in red font. The molecular weight marker lane on the far left is marked by ‘M’. C) Negative stain electron microscopy of fraction 4. Some of the F-ATP synthase complexes that appear to be tetrameric F-ATP synthase are encircled in red. Scale bar 50 nm.

**Figure 5:**
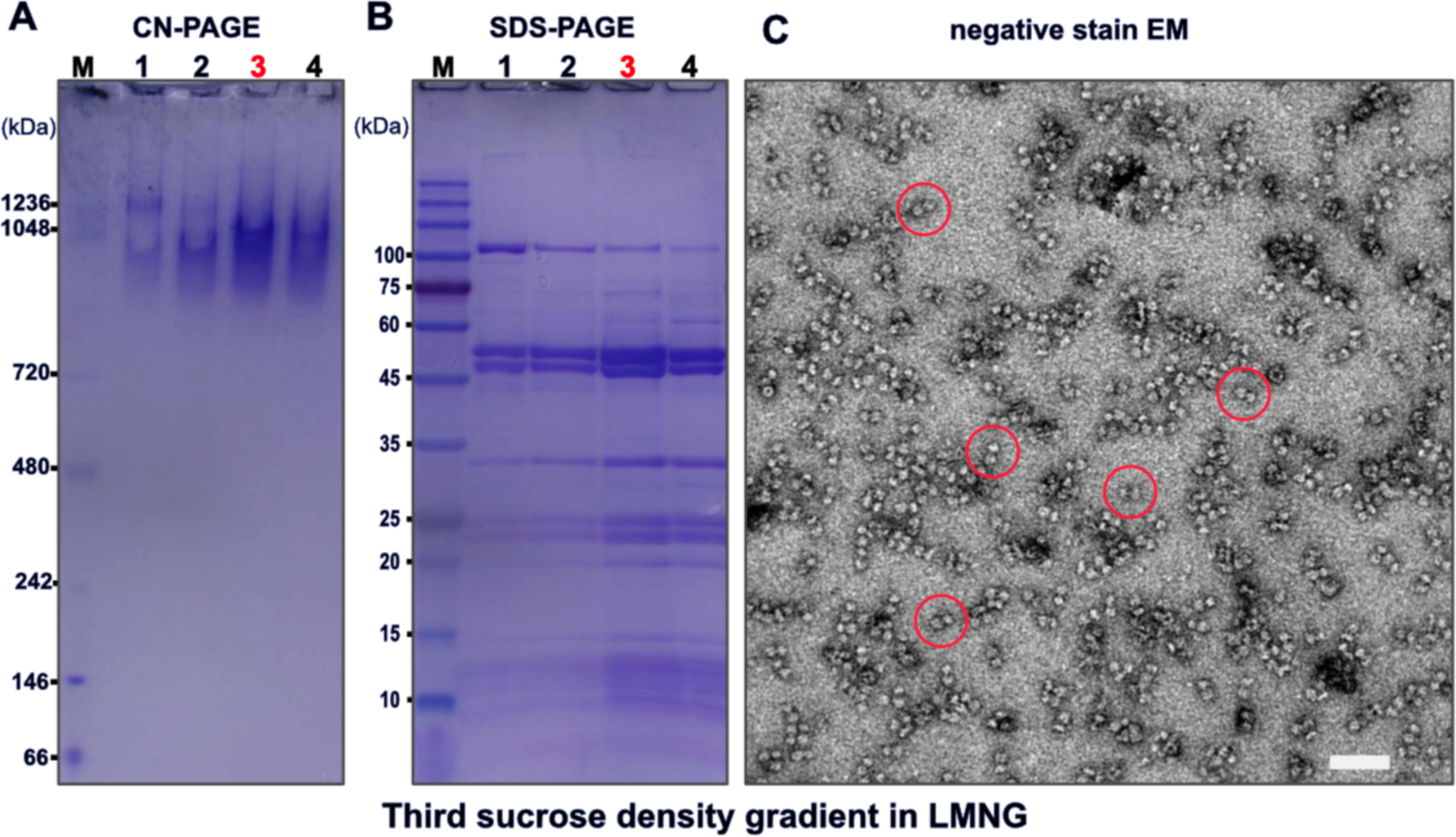
Analysis of the third continuous sucrose LMNG density gradient fractions by native gel electrophoresis, denaturing gel electrophoresis and negative stain electron microscopy. A) Clear native gel electrophoresis of fractions collected from bottom to top of the gradient with the fraction enriched in monomeric F-ATP synthase and used for negative stain electron microscopy marked in red font. The molecular weight marker lane on the far left is marked by ‘M’. B) Denaturing gel electrophoresis of fractions collected from bottom to top of the gradient as analyzed in A). The fraction enriched in monomeric F-ATP synthase and used for negative stain electron microscopy in marked in red font. Molecular weight marker are indicated on the left side of the gel. C) Negative stain electron microscopy of fraction 3. Some of the F-ATP synthase complexes that appear to be monomeric F-ATP synthase are encircled in red. Scale bar 50 nm.

**Figure 6:**
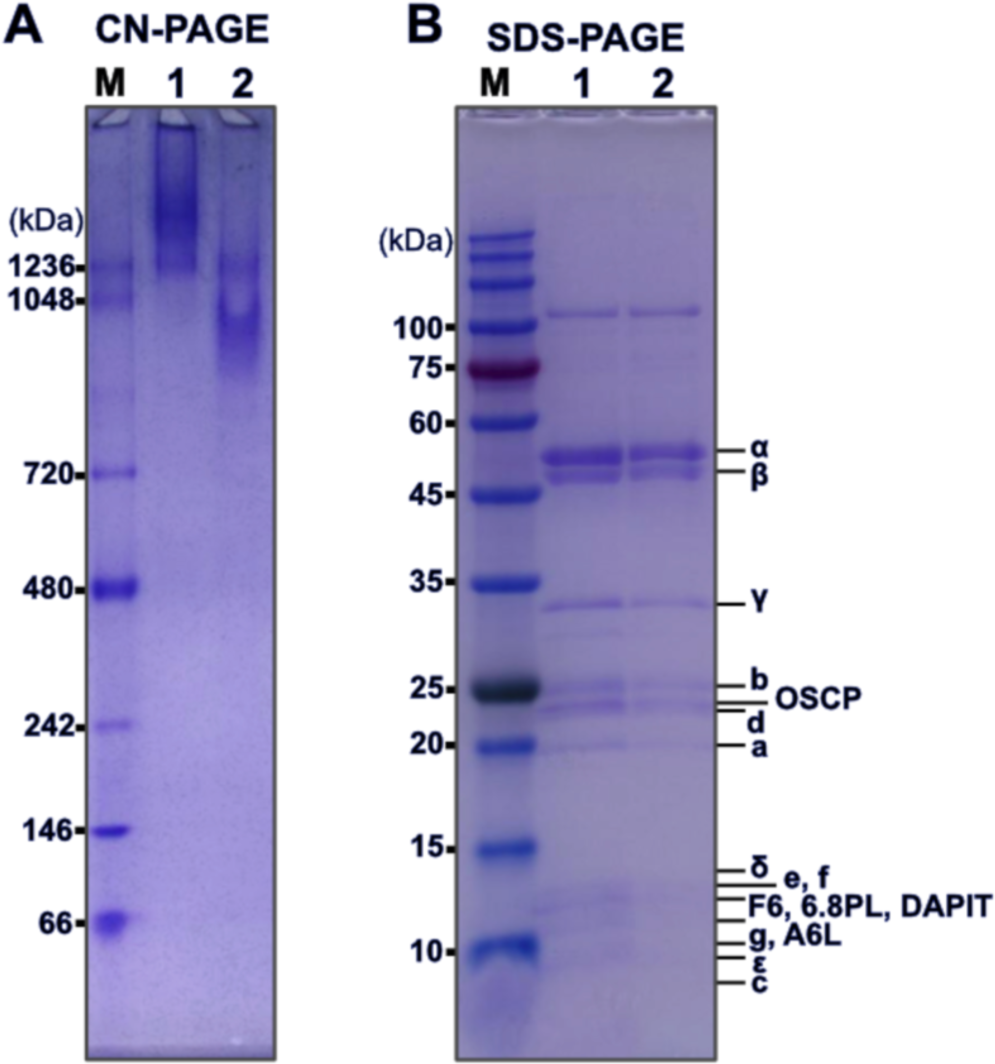
Analysis of the final purified tetramer and monomer F-ATP synthase by native gel electrophoresis and denaturing gel electrophoresis after storage for 10 days. A) Clear native gel electrophoresis of the final tetramer preparation in Lane 1 and final monomer preparation in Lane 2. The molecular weight marker lane on the far left is marked by ‘M’. B) Denaturing gel electrophoresis of the final tetramer preparation in Lane 1 and final monomer preparation in Lane 2. The molecular weight marker lane on the far left is marked by ‘M’.

## Discussion

The aim of protein purification is to isolate the target complex from its surrounding cellular components in order to allow its in vitro investigation in the absence of interference and as a result obtain experimental data that is sufficiently interpretable to expand our understanding of the target complex biology. For a successful implementation of this reductionist approach to molecular biology ideally the target complex is purified from its cellular environment to homogeneity without compromising its structural and functional integrity. In the case of mammalian F-ATP synthase this is a formidable task as a consequence of its complex and fragile multisubunit composition, which is known to be very sensitive to the disruptive effect of conventional detergents, presumeably due to the loss of bound lipids(50). A further complication stems from the circumstance that F-ATP synthase is a rotary motor nano-machine that is sensitive to mechanical stress. Furthermore, the dynamic oligomeric state of mammalian F-ATP synthase between higher oligomer, tetramers, dimers and monomers(51) complicates the purification of the complex in an oligomeric pure state. However, conclusive in vitro experiments that are able to answer important questions on higher functions mediated by the supernumerary subunits and the oligomeric state clearly necessitate purification procedures that preserve functional and structural integrity for all oligomeric states of interest. Traditionally the enrichment of an enzyme during purification is monitored by increasing its specific activity. Since we aimed at the isolation of the IF1 bound oligomer, the opposite had to be achieved: minimized enzymatic activity with maximized band intensity in denaturing and native electrophoresis. Here, we have shown that under buffer conditions that favor IF1 binding and oligomer stabilization it is possible to obtain large amounts of pure and stable F-ATP synthase from bovine heart mitochondria without the use of chromatography steps or the use of affinity resins. The key for the here described purification strategy is the separation of very large higher oligomers of IF1 bound F-ATP synthase from other components of the respiratory chain after detergent mediated solubilization of the inner mitochondrial membrane. A second sucrose step density gradient is then sufficient to obtain very pure, however, oligomeric state mixes F-ATP synthase. We demonstrate that enrichment of two oligomeric states, namely tetramer and monomer, is feasible when using a third continuous sucrose density gradient. Since the tetrameric form of F-ATP synthase represents the smallest higher oligomer, a dimer of dimers, this opens the way to conduct in vitro experiments probing the functional meaning of oligomerization for the mammalian complex. Considering that questions in regard to how the oligomeric state is affecting the F-ATP synthase core function of ATP synthesis remain unanswered, this is of interest. Furthermore, open questions in regard to monomeric F-ATP synthase ability to elicit conductance states of the PTP might be more easy to investigate when making use of the here reported isolation procedures. The here described approach of isolating monomeric F-ATP synthase is unique in that the immediate input to the monomer gradient is IF1 bound oligomeric F-ATP synthase. As a consequence the presence of mixed levels of structural integrity is avoided and a high level of structural homogeneity provided. Another point of interest is whether oligomerization of mitochondrial F-ATP synthase requires a membrane or not(52). Oligomer contacts might involve complex bound lipids that are possibly stripped off during chromatography steps, potentially limiting the ability to reconstitute oligomeric F-ATP synthase in solution. If the presence of a membrane is not essential, we expect that in solution oligomerization might be observable for column-free purified mitochondrial F-ATP synthase. Our current structural understanding of mammalian F-ATP synthase is based on reported single particle cryo-EM structures of the porcine tetramer(11), a monomer map obtained from the bovine dimer(10), the ovine monomer(9) and the human monomer(35). Although important achievements, their resolution and map resolvability is still wanting in all structures. If these limitations are at least partially the result of compositional heterogeneity, the here described column-free purification approach might be useful to improve resolution, map quality and atomic models for both tetrameric and monomeric F-ATP synthase isolated from mammalian heart muscle tissue or human cell cultures.

An obvious drawback of our purification approach is the large amount of muscle tissue needed. Though the here used total amount of 600 g muscle tissue for one purification is relatively modest, study of human and mutant F-ATP synthase from cell cultures will require the adjustment of the current protocol to smaller amounts of starting material. Tetrameric IF1 bound F-ATP synthase as reported for the porcine heart muscle mitochondria and enriched in our GDN based third continuous sucrose density gradient, likely forms when after animal sacrifice the lack of oxygen in heart muscle cells causes a breakdown of the pmf and rapid acidification causes binding of two dimeric IF1 to neighboring F-ATP synthase dimers. We succeeded to enrich tetrameric F-ATP synthase to a level that appears to be sufficient for structure determination by single particle cryo-EM. However, both for structural and also functional studies enrichment to 100% is desirable and should be the aim of further improvements of the current protocol. The purification procedure described here relies on the balanced use of the two novel detergents GDN and LMNG. Their relative amount can be used to shift the oligomeric state in solution without affecting the integrity of the monomeric complex. Further variants of these two, lipid like detergents might ease this in solution manipulation of various oligomeric states and perhaps even allow for the efficient auto-insertion of higher oligomers, which unfortunately is not supported by the presence of GDN as the stabilizing detergent(38).

In recent years F-ATP synthase has attracted attention as a therpeutic drug target(53, 54). Indeed, the development of (+)-Epicatechin as a small molecule mimic of IF1 demonstrates that human F-ATP synthase could be crucial molecular player for drug development aimed at mitochondrial disorders(17). As such, our newly developed large-scale column-free purification might evolve into a cornerstone for mitochondrial F-ATP synthase targeted drug screening, especially, if screens are aimed at finding pharmacological compounds that target oligomeric F-ATP synthase. The here reported purification strategy will hopefully be explored, implemented and improved by a wide range of investigators interested in the biology of mammalian F-ATP synthase and thus contribute to our understanding of its structure and function.

## Data availability

All data is available upon reasonable request to the corresponding authors.

## Acknowledgements

## Acknowledgements

We would like to thank Kyoko Shinzawa-Itoh, Shinya Yoshikawa and Genji Kurisu for their kind support for this project.

## Author contributions

Chimari Jiko: Conceptualization, Methodology, Validation, Investigation, Resources, Writing - Review&Editing, Funding acquisition Yukio Morimoto: Resources, Writing - Review&Editing Tomitake Tsukihara: Resources, Writing - Review&Editing, Supervision Christoph Gerle: Conceptualization, Methodology, Validation, Resources, Writing-Original Draft, Writing - Review&Editing, Funding acquisition

## Conflict of interested

A patent application has been submitted by C.J., Y.M., C.G. and Kyoto University based on the results of this study.

## Supporting Information

**Supporting Figure 1:**
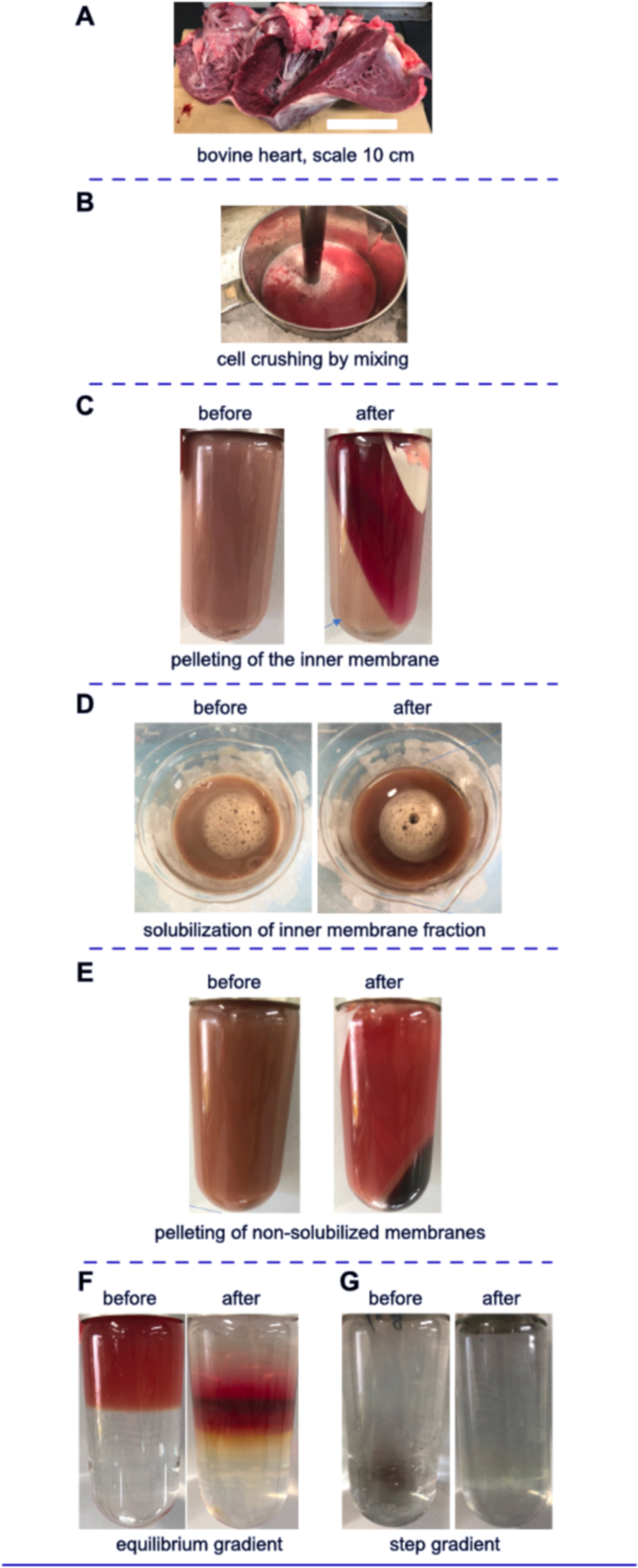
Gallery of experimental steps used to go from bovine heart muscle tissue to colorless F-ATP synthase oligomer enriched sucrose density gradient fraction. A) Typical bovine heart used for F-ATP synthase purification. B) Muscle cell crushing by Polytron mixing. C) Mitochondrial inner membrane pelleting by ultracentrifugation. Membrane fraction is indicated by a blue arrow on the lower left in ‘after’ and residual oil can be seen as a white-yellow substance on the upper right in ‘after’. D) Detergent driven mitochondrial inner membrane solubilization while stirring on ice. E) Pelleting of non-solubilized material by ultracentrifugation. F) The first sucrose density gradient ultracentrifugation by equilibrium centrifugation for separation of IF1 bound higher oligomer F-ATP synthase from other components of the respiratory chain. Left side: after loading; right side: after centrifugation. G) The second density gradient ultracentrifugation using a step gradient for further removal of contaminants. Left side: after loading; right side: after centrifugation.

**Supporting Figure 2:**
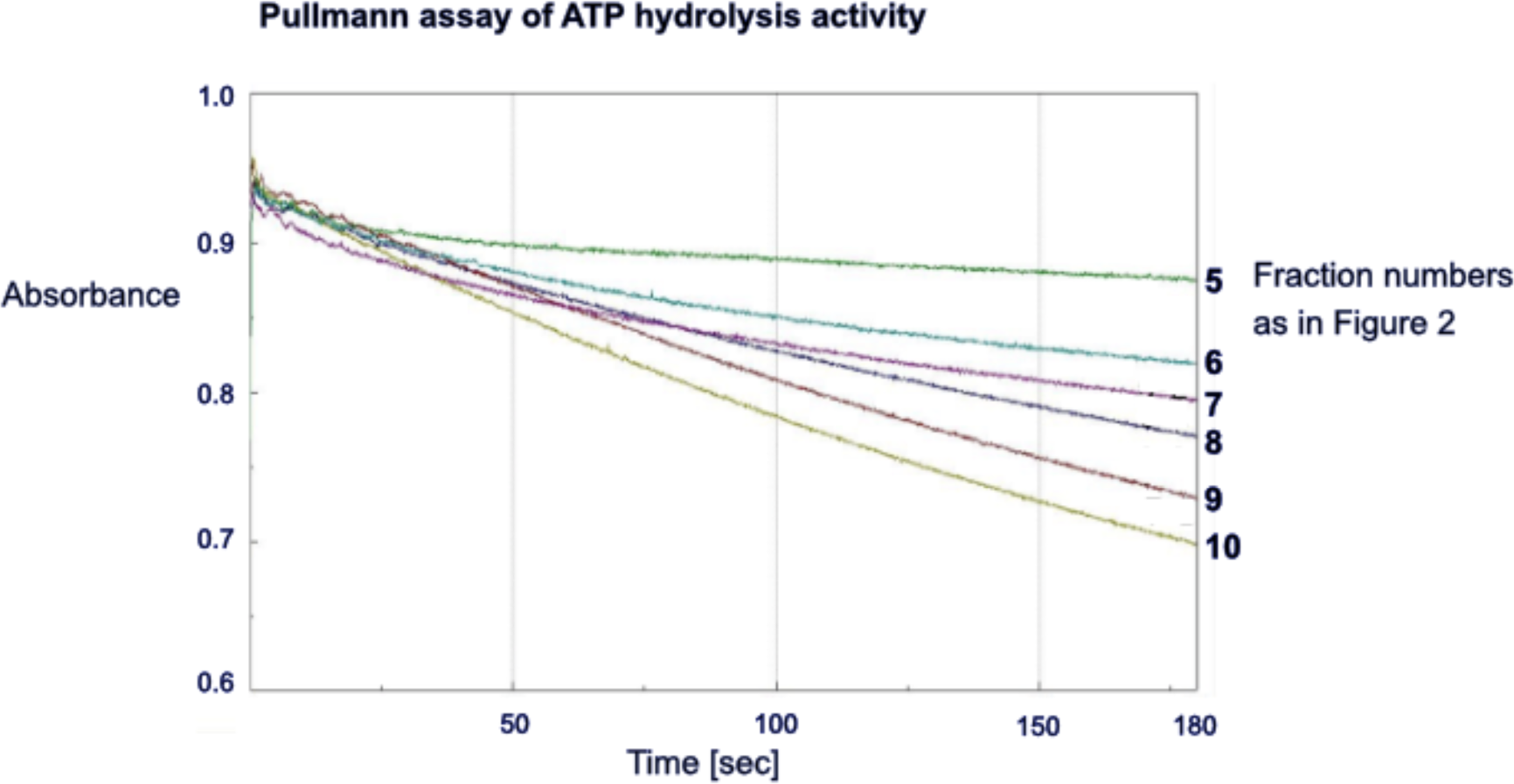
ATP hydrolase activity of fractions collected after the first sucrose density ultracentrifugation. Fractions 5 and 6 exhibit very little ATPase activity indicating the presence of IF1 bound F-ATP synthase.

**Supporting Figure 3:**
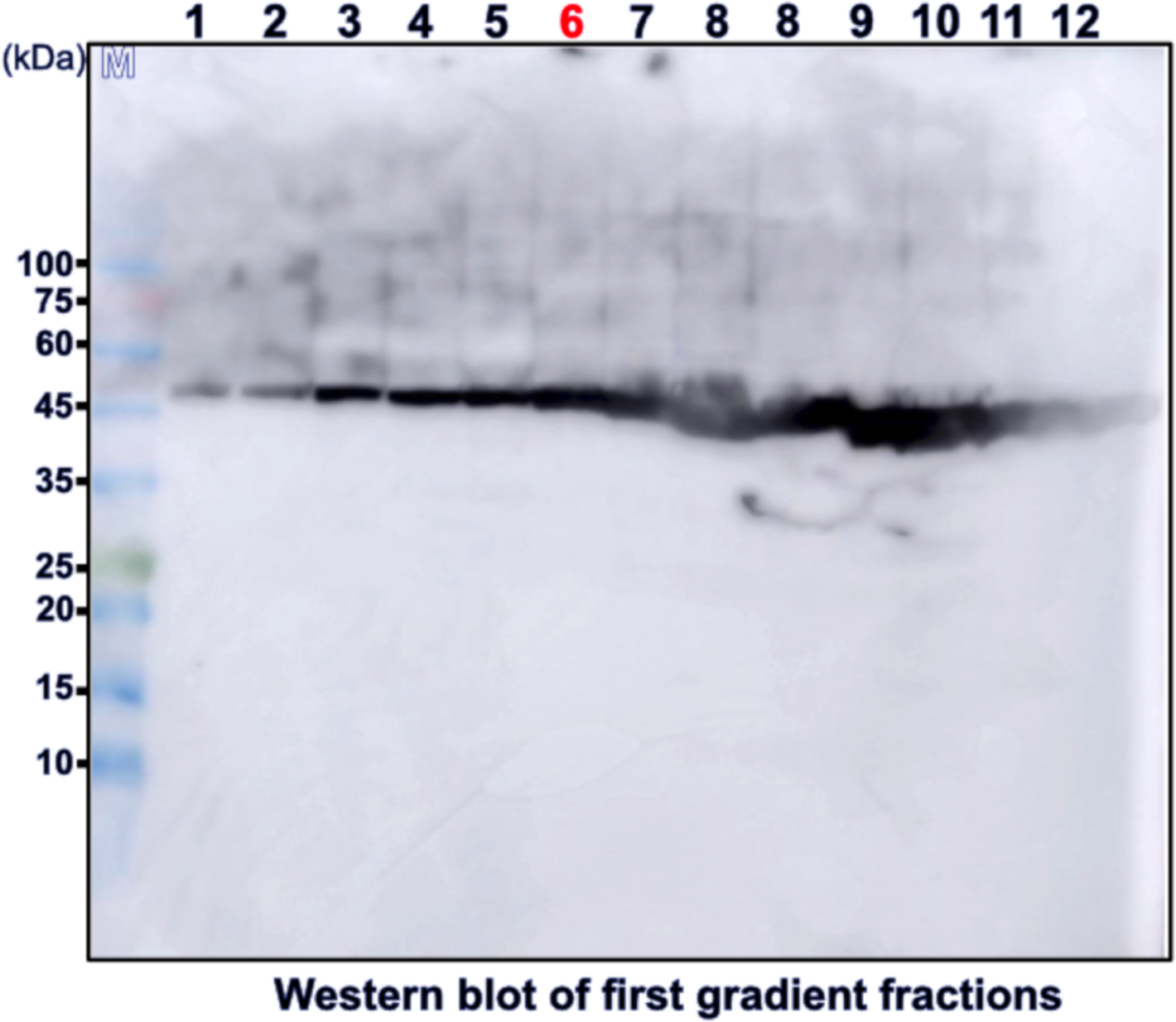
Western blot against F-ATP synthase β subunit of the first gradient fractions. Fractions used are identical to those of Figure 2. The β subunit is clearly detected in all fractions of the gradient including the lower, almost colorless fractions.

## References

1. Yoshida, M., Muneyuki, E., and Hisabori, T. (2001) ATP synthase--a marvellous rotary engine of the cell. Nat. Rev. Mol. cell Biol. 2, 669–677

2. Kühlbrandt, W. (2019) Structure and Mechanisms of F-Type ATP Synthases. Annu. Rev. Biochem. 10.1146/annurev-biochem-013118-110903

3. Walker, J. E. (2013) The ATP synthase: the understood, the uncertain and the unknown. Biochem. Soc. Trans. 41, 1–16

4. Carter, A. P., Mühleip, A., Mccomas, S. E., and Amunts, A. (2019) Structure of a mitochondrial ATP synthase with bound native cardiolipin. 10.7554/eLife.51179

5. Mühleip, A., Kock Flygaard, R., Ovciarikova, J., Lacombe, A., Fernandes, P., Sheiner, L., and Amunts, A. (2021) ATP synthase hexamer assemblies shape cristae of Toxoplasma mitochondria. Nat. Commun. 10.1038/s41467-020-20381-z

6. Flygaard, R. K., Mühleip, A., Tobiasson, V., and Amunts, A. (2020) Type III ATP synthase is a symmetry-deviated dimer that induces membrane curvature through tetramerization. Nat. Commun. 10.1038/s41467-020-18993-6

7. Hahn, A., Vonck, J., Mills, D. J., Meier, T., and Kühlbrandt, W. (2018) Structure, mechanism, and regulation of the chloroplast ATP synthase. Science (80-.). 10.1126/SCIENCE.AAT4318/SUPPL_FILE/AAT4318_HAHN_SM.PDF

8. A. Sobti, M., Smits, C., Wong, A. S. W., Ishmukhametov, R., Stock, D., Sandin S., and Stewart, G. (2016) Cryo-EM structures of the autoinhibited E. coli ATP synthase in three rotational states. Elife. 10.7554/ELIFE.21598

9. Pinke, G., Zhou, L., and Sazanov, L. A. (2020) Cryo-EM structure of the entire mammalian F-type ATP synthase. Nat. Struct. Mol. Biol. 10.1038/s41594-020-0503-8

10. Spikes, T. E., Montgomery, M. G., and Walker, J. E. (2020) Structure of the dimeric ATP synthase from bovine mitochondria. Proc. Natl. Acad. Sci. U. S. A. 10.1073/pnas.2013998117

11. Gu, J., Zhang, L., Zong, S., Guo, R., Liu, T., Yi, J., Wang, P., Zhuo, W., and Yang, M. (2019) Cryo-EM structure of the mammalian ATP synthase tetramer bound with inhibitory protein IF1. Science (80-.). 10.1126/science.aaw4852

12. Guo, H., Bueler, S. A., and Rubinstein, J. L. (2017) Atomic model for the dimeric FO region of mitochondrial ATP synthase. Science (80-.). 358, 936–940

13. Pullman, M. E., and Monroy, G. C. (1963) A Naturally Occurring Inhibitor of Mitochondrial Adenosine Triphosphatase. J. Biol. Chem. 238, 3762–3769

14. García, J. J., Morales-Ríos, E., Cortés-Hernández, P., and Rodríguez-Zavala, J. S. (2006) The inhibitor protein (IF1) promotes dimerization of the mitochondrial F1F0-ATP synthase. Biochemistry. 45, 12695–12703

15. Minauro-Sanmiguel, F., Wilkens, S., and Garcia, J. J. (2005) Structure of dimeric mitochondrial ATP synthase: Novel F0 bridging features and the structural basis of mitochodrial cristae biogenesis. Proc. Natl. Acad. Sci. U. S. A. 102, 12356–12358

16. Campanella, M., Casswell, E., Chong, S., Farah, Z., Wieckowski, M. R., Abramov, A. Y., Tinker, A., and Duchen, M. R. (2008) Regulation of Mitochondrial Structure and Function by the F1Fo-ATPase Inhibitor Protein, IF1. Cell Metab. 8, 13–25

17. Acin-Perez, R., Beninc, C., Fernandez Del Rio, L., Shu, C., Baghdasarian, S., Zanette, V., Gerle, C., Jiko, C., Khairallah, R., Khan, S., Rincon, D., Pacheco, F., Shabane, B., Erion, K., Masand, R., Dugar, S., Ghenoiu, C., Schreiner, G., Stiles, L., Liesa, M., and Shirihai, O. S. (2023) Inhibition of ATP synthase reverse activity restores energy homeostasis in mitochondrial pathologies. EMBO J. 42, e111699

18. Giorgio, V., von Stockum, S., Antoniel, M., Fabbro, A., Fogolari, F., Forte, M., Glick, G. D., Petronilli, V., Zoratti, M., Szabó, I., Lippe, G., and Bernardi, P. (2013) Dimers of mitochondrial ATP synthase form the permeability transition pore. Proc. Natl. Acad. Sci. U. S. A. 110, 5887–5892

19. Urbani, A., Giorgio, V., Carrer, A., Franchin, C., Arrigoni, G., Jiko, C., Abe, K., Maeda, S., Shinzawa-Itoh, K., Bogers, J. F. M., McMillan, D. G. G., Gerle, C., Szabò, I., and Bernardi, P. (2019) Purified F-ATP synthase forms a Ca2+-dependent high-conductance channel matching the mitochondrial permeability transition pore. Nat. Commun. 10.1038/s41467-019-12331-1

20. Gerle, C. (2020) Mitochondrial F-ATP synthase as the permeability transition pore. Pharmacol. Res. 160, 105081

21. Pullman, M. E., Penefsky, H. S., Datta, A., and Racker, E. (1960) Partial resolution of the enzymes catalyzing oxidative phosphorylation. I. Purification and properties of soluble dinitrophenol-stimulated adenosine triphosphatase. J. Biol. Chem. 235, 3322– 3329

22. Kagawa, Y., and Racker, E. (1966) Partial resolution of the enzymes catalyzing oxidative phosphorylation. X. Correlation of morphology and function in submitochondrial particles. J. Biol. Chem. 241, 2475–2482

23. Lutter, R., Saraste, M., Van Walraven, H. S., Runswick, M. J., Finel, M., Deatherage, J. F., and Walker, J. E. (1993) F1F0-ATP synthase from bovine heart mitochondria: Development of the purification of a monodisperse oligomycin-sensitive ATPase. Biochem. J. 10.1042/bj2950799

24. Buchanan, S. K., and Walker, J. E. (1996) Large-scale chromatographic purification of F1F0-ATPase and complex I from bovine heart mitochondria. Biochem. J. 318 (Pt 1, 343– 349

25. Wilkens, S., Inoue, T., and Forgac, M. (2004) Three-dimensional structure of the vacuolar ATPase. Localization of subunit H by difference imaging and chemical cross-linking. J. Biol. Chem. 279, 41942–41949

26. Maeda, S., Shinzawa-Itoh, K., Mieda, K., Yamamoto, M., Nakashima, Y., Ogasawara, Y., Jiko, C., Tani, K., Miyazawa, A., Gerle, C., and Yoshikawa, S. (2013) Two-dimensional crystallization of intact F-ATP synthase isolated from bovine heart mitochondria. *Acta Crystallogr. Sect. F*, Struct. Biol. Cryst. Commun. 69, 1368–1370

27. Jiko, C., Davies, K. M., Shinzawa-Itoh, K., Tani, K., Maeda, S., Mills, D. J., Tsukihara, T., Fujiyoshi, Y., Uhlbrandt, W., and Gerle, C. Bovine F 1 F o ATP synthase monomers bend the lipid bilayer in 2D membrane crystals. 10.7554/eLife.06119.001

28. Shimada, S., Maeda, S., Hikita, M., Mieda-Higa, K., Uene, S., Nariai, Y., and Shinzawa-Itoh, K. (2018) Solubilization conditions for bovine heart mitochondrial membranes allow selective purification of large quantities of respiratory complexes I, III, and V. Protein Expr. Purif. 150, 33–43

29. Runswick, M. J., Bason, J. V, Montgomery, M. G., Robinson, G. C., Fearnley, I. M., and Walker, J. E. (2013) The affinity purification and characterization of ATP synthase complexes from mitochondria. Open Biol. 3, 120160

30. Lutter, R., Abrahams, J. P., van Raaij, M. J., Todd, R. J., Lundqvist, T., Buchanan, S. K., Leslie, A. G., and Walker, J. E. (1993) Crystallization of F1-ATPase from bovine heart mitochondria. J. Mol. Biol. 229, 787–790

31. Abrahams, J. P., Leslie, A. G., Lutter, R., and Walker, J. E. (1994) Structure at 2.8 A resolution of F1-ATPase from bovine heart mitochondria. Nature. 370, 621–628

32. Chae, P. S., Rasmussen, S. G. F., Rana, R. R., Gotfryd, K., Kruse, A. C., Manglik, A., Cho, K. H., Nurva, S., Gether, U., Guan, L., Loland, C. J., Byrne, B., Kobilka, B. K., and Gellman, S. H. (2012) A new class of amphiphiles bearing rigid hydrophobic groups for solubilization and stabilization of membrane proteins. Chem. - A Eur. J. 18, 9485–9490

33. Chae, P. S., Rasmussen, S. G., Rana, R. R., Gotfryd, K., Chandra, R., Goren, M. A., Kruse, A. C., Nurva, S., Loland, C. J., Pierre, Y., Drew, D., Popot, J. L., Picot, D., Fox, B. G., Guan, L., Gether, U., Byrne, B., Kobilka, B., and Gellman, S. H. (2010) Maltose-neopentyl glycol (MNG) amphiphiles for solubilization, stabilization and crystallization of membrane proteins. Nat. Methods. 7, 1003–1008

34. Hauer, F., Gerle, C., Fischer, N., Oshima, A., Shinzawa-Itoh, K., Shimada, S., Yokoyama, K., Fujiyoshi, Y., and Stark, H. (2015) GraDeR: Membrane Protein Complex Preparation for Single-Particle Cryo-EM. Structure. 10.1016/j.str.2015.06.029

35. Lai, Y., Zhang, Y., Zhou, S., Xu, J., Du, Z., Feng, Z., Yu, L., Zhao, Z., Wang, W., Tang, Y., Yang, X., Guddat, L. W., Liu, F., Gao, Y., Rao, Z., and Gong, H. (2023) Structure of the human ATP synthase. Mol. Cell. 10.1016/J.MOLCEL.2023.04.029

36. Cannino, G., Urbani, A., Gaspari, M., Varano, M., Negro, A., Filippi, A., Ciscato, F., Masgras, I., Gerle, C., Tibaldi, E., Brunati, A. M., Colombo, G., Lippe, G., Bernardi, P., and Rasola, A. (2022) The mitochondrial chaperone TRAP1 regulates F-ATP synthase channel formation. Cell Death Differ. 2022 2912. 29, 2335–2346

37. Quintana-Cabrera, R., Quirin, C., Glytsou, C., Corrado, M., Urbani, A., Pellattiero, A., Calvo, E., Vázquez, J., Enríquez, J. A., Gerle, C., Soriano, M. E., Bernardi, P., and Scorrano, L. (2018) The cristae modulator Optic atrophy 1 requires mitochondrial ATP synthase oligomers to safeguard mitochondrial function. Nat. Commun. 2018 91. 9, 1–13

38. Godoy-Hernandez, A., Asseri, A. H., Purugganan, A. J., Jiko, C., de Ram, C., Lill, H., Pabst, M., Mitsuoka, K., Gerle, C., Bald, D., and McMillan, D. G. G. (2022) Rapid and Highly Stable Membrane Reconstitution by LAiR Enables the Study of Physiological Integral Membrane Protein Functions. ACS Cent. Sci. 9, 3, 494–507

39. Pinke, G., Zhou, L., and Sazanov, L. A. (2020) Cryo-EM structure of the entire mammalian F-type ATP synthase. Nat. Struct. Mol. Biol. 27, 1077–1085

40. Lai, Y., Zhang, Y., Zhou, S., Xu, J., Du, Z., Feng, Z., Yu, L., Zhao, Z., Wang, W., Tang, Y., Yang, X., Guddat, L. W., Liu, F., Gao, Y., Rao, Z., and Gong, H. (2023) Short article Structure of the human ATP synthase Short article Structure of the human ATP synthase. Mol. Cell. 10.1016/j.molcel.2023.04.029

41. A. Mnatsakanyan, N., Llaguno, M. C., Yang, Y., Yan, Y., Weber, J., Sigworth F. J., and Jonas, E. (2019) A mitochondrial megachannel resides in monomeric F1FO ATP synthase. Nat. Commun. 10.1038/s41467-019-13766-2

42. Tsukihara, T., Aoyama, H., Yamashita, E., Tomizaki, T., Yamaguchi, H., Shinzawa-Itoh, K., Nakashima, R., Yaono, R., and Yoshikawa, S. (1996) The whole structure of the 13-subunit oxidized cytochrome c oxidase at 2.8 A. Sci. (New York, NY). 272, 1136–1144

43. Khatter, H., Myasnikov, A. G., Mastio, L., Billas, I. M. L., Birck, C., Stella, S., and Klaholz, B. P. Purification, characterization and crystallization of the human 80S ribosome. 10.1093/nar/gkt1404

44. Letts, J. A., Fiedorczuk, K., and Sazanov, L. A. (2016) The architecture of respiratory supercomplexes. Nat. 2016 5377622. 537, 644–648

45. Pan, X., Ma, J., Su, X., Cao, P., Chang, W., Liu, Z., Zhang, X., and Li, M. (2018) Structure of the maize photosystem I supercomplex with light-harvesting complexes I and II. Science (80-.). 360, 1109–1113

46. Mochizuki, M., Aoyama, H., Shinzawa-Itoh, K., Usui, T., Tsukihara, T., and Yoshikawa, S. (1999) Quantitative reevaluation of the redox active sites of crystalline bovine heart cytochrome c oxidase. J. Biol. Chem. 274, 33403–33411

47. Chorev, D. S., Baker, L. A., Wu, D., Beilsten-Edmands, V., Rouse, S. L., Zeev-Ben-Mordehai, T., Jiko, C., Samsudin, F., Gerle, C., Khalid, S., Stewart, A. G., Matthews, S. J., Grünewald, K., and Robinson, C. V. (2018) Protein assemblies ejected directly from native membranes yield complexes for mass spectrometry. Science (80-.). 362, 829–834

48. BELTRÁN, C., de GÓMEZ-PUYOU, M. T., GÓMEZ-PUYOU, A., and Darszon, A. (1984) Release of the inhibitory action of the natural ATPase inhibitor protein on the mitochondrial ATPase. Eur. J. Biochem. 144, 151–157

49. Wittig, I., and Schägger, H. (2005) Advantages and limitations of clear-native PAGE. Proteomics. 5, 4338–4346

50. Meyer, B., Wittig, I., Trifilieff, E., Karas, M., and Schägger, H. (2007) Identification of two proteins associated with mammalian ATP synthase. Mol. Cell. Proteomics. 10.1074/mcp.M700097-MCP200

51. Ader, N. R., Hoffmann, P. C., Ganeva, I., Borgeaud, A. C., Wang, C., Youle, R. J., and Kukulski, W. (2019) Molecular and topological reorganizations in mitochondrial architecture interplay during bax-mediated steps of apoptosis. Elife. 10.7554/ELIFE.40712

52. Davies, K. M., Anselmi, C., Wittig, I., Faraldo-Gómez, J. D., and Kühlbrandt, W. (2012) Structure of the yeast F1Fo-ATP synthase dimer and its role in shaping the mitochondrial cristae. Proc. Natl. Acad. Sci. U. S. A. 109, 13602–13607

53. Dautant, A., Meier, T., Hahn, A., Tribouillard-Tanvier, D., di Rago, J. P., and Kucharczyk, R. (2018) ATP Synthase Diseases Of Mitochondrial Genetic Origin. Front. Physiol. 9, 329

54. Vlasov, A. V., Osipov, S. D., Bondarev, N. A., Uversky, V. N., Borshchevskiy, V. I., Yanyushin, M. F., Manukhov, I. V., Rogachev, A. V., Vlasova, A. D., Ilyinsky, N. S., Kuklin, A. I., Dencher, N. A., and Gordeliy, V. I. (2022) ATP synthase FOF1 structure, function, and structure-based drug design. Cell. Mol. Life Sci. 79, 1–27

