## Supplemental Figures for "Large-scale column-free purification of bovine F-ATP synthase"

### Supporting Information

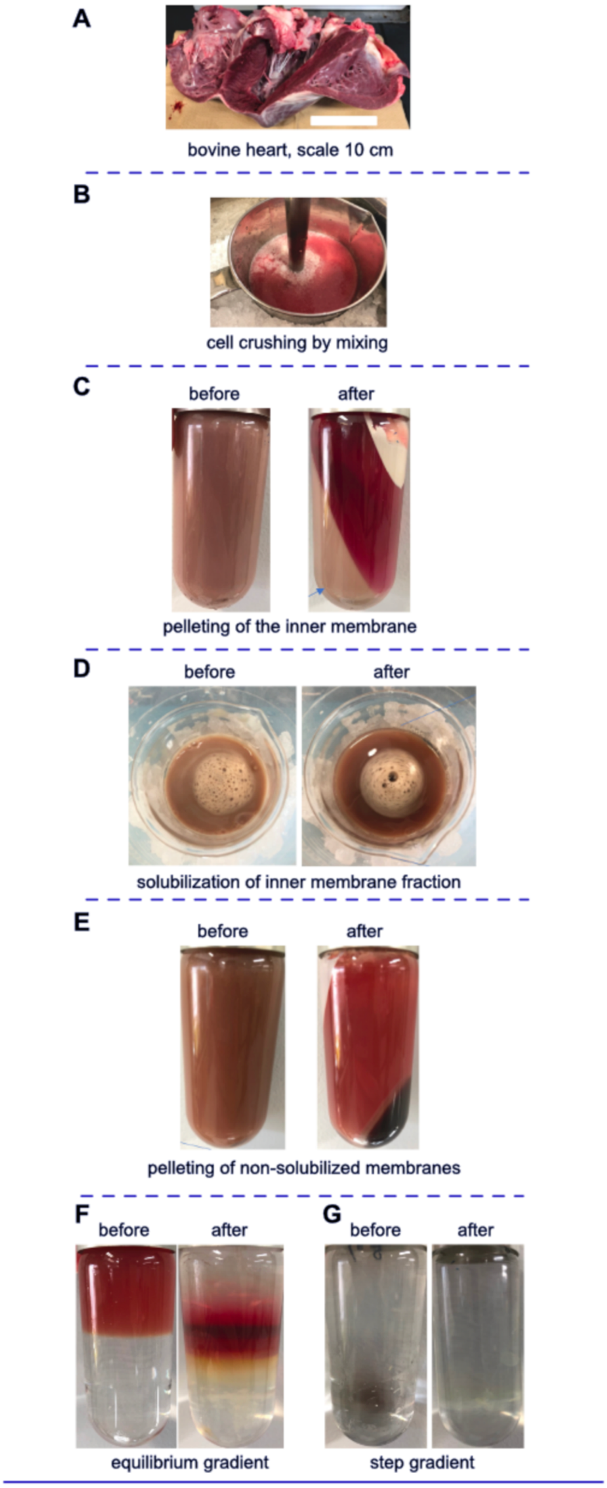

**Supporting Figure 1: Gallery of experimental steps used to go from bovine heart muscle tissue to colorless F-ATP synthase oligomer enriched sucrose density gradient fraction.** A) Typical bovine heart used for F-ATP synthase purification. B) Muscle cell crushing by Polytron mixing. C) Mitochondrial inner membrane pelleting by ultracentrifugation. Membrane fraction is indicated by a blue arrow on the lower left in 'after' and residual oil can be seen as a white-yellow substance on the upper right in 'after'. D) Detergent driven mitochondrial inner membrane solubilization while stirring on ice. E) Pelleting of non-solubilized material by ultracentrifugation. F) The first sucrose density gradient ultracentrifugation by equilibrium centrifugation for separation of IF1 bound higher oligomer F-ATP synthase from other components of the respiratory chain. Left side: after loading; right side: after centrifugation. G) The second density gradient ultracentrifugation using a step gradient for further removal of contaminants. Left side: after loading; right side: after centrifugation.

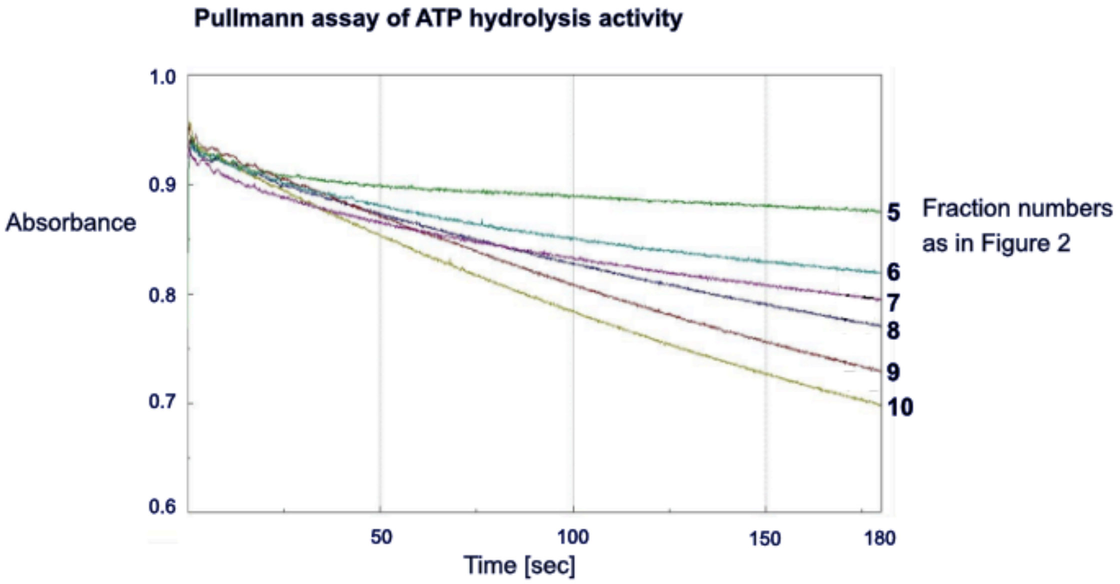


**Supporting Figure 2: ATP hydrolase activity of fractions collected after the first sucrose density ultracentrifugation.** Fractions 5 and 6 exhibit very little ATPase activity indicating the presence of IF1 bound F-ATP synthase.

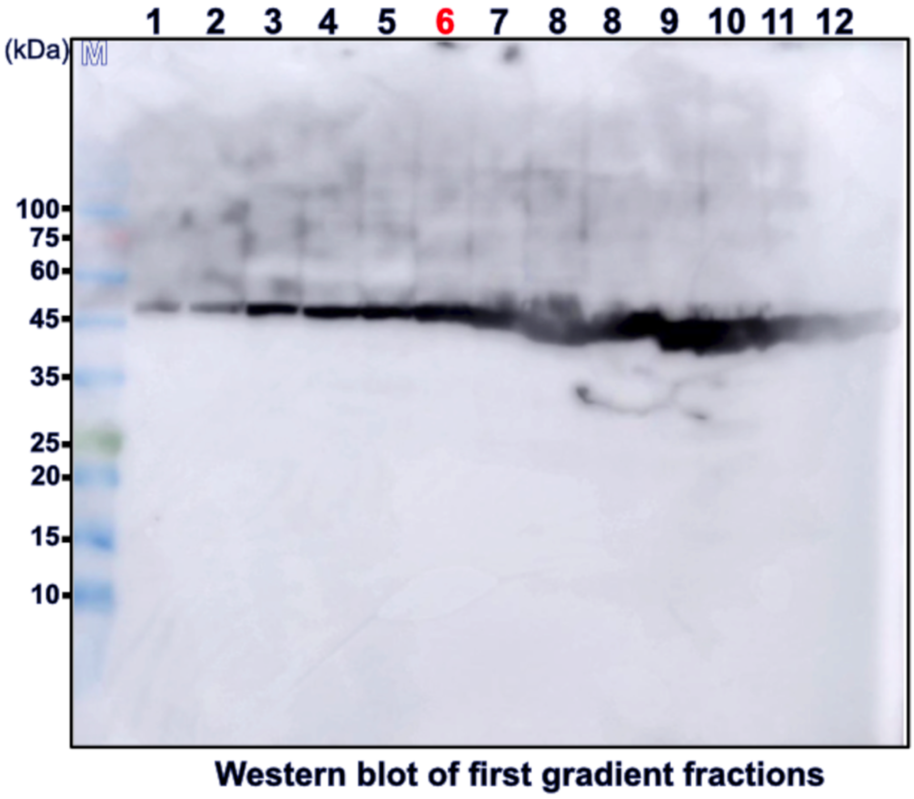


**Supporting Figure 3: Western blot against F-ATP synthase β subunit of the first gradient fractions.** Fractions used are identical to those of Figure 2. The β subunit is clearly detected in all fractions of the gradient including the lower, almost colorless fractions.
